# Circadian fluctuations in glucocorticoid level predict perceptual discrimination sensitivity

**DOI:** 10.1101/2020.10.07.330282

**Authors:** Jonas Obleser, Jens Kreitewolf, Ricarda Vielhauer, Fanny Lindner, Carolin David, Henrik Oster, Sarah Tune

## Abstract

Slow neurobiological rhythms, such as the circadian secretion of glucocorticoid (GC) hormones, modulate a wide variety of body functions. Whether and how such endocrine fluctuations also exert an influence on perceptual abilities is largely uncharted. Here, we show that phasic, moderate increases in GC availability prove beneficial to auditory discrimination. In an age-varying sample of N = 68 healthy human participants, we characterise the covariation of saliva cortisol with perceptual sensitivity in an auditory pitch-discrimination task at five time points across the sleep--wake cycle. First, momentary saliva cortisol levels were captured well by the time relative to wake-up and overall sleep duration. Second, within individuals, higher cortisol level just prior to behavioural testing predicted better pitch discrimination ability, expressed as a steepened psychometric curve. This effect of glucocorticoids held under a set of statistical control models. Our results pave the way for more in-depth studies on neuroendocrinological determinants of sensory encoding and perception.

## Introduction

Most physiological functions in humans exert circadian rhythmicity (Dibner et al., 2010). That is, bodily homeostatic functions oscillate with a period of about 24 hours and are vital in adapting the organism to its environment (Spiga et al., 2014). These functions are regulated through the endogenous circadian clock system with the suprachiasmatic nucleus (SCN) as pacemaker, synchronising subordinate tissue clocks located throughout the body (Dibner et al., 2010).

While sensory, perceptual, and cognitive functions all have been shown to also be subject to—much faster—rhythmicity and to covary with brain states at the sub-second (“neural oscillations”, e.g. Henry and Obleser, 2012) or seconds-to-minutes scale (e.g., Park et al., 2014; Rebollo et al., 2018), a potential circadian role of the endocrine system in the regulation of sensation, and perception in particular, has received much less attention.

The endocrine system, with glucocorticoids (GC) as the main effector of the hypothalamic-pituitary-adrenal (HPA) axis, exhibits also prominent circadian rhythmicity (Spencer and Deak, 2017). Cortisol as the major human endogenous glucocorticoid is recognised in psychophysiological research primarily as a stress hormone, reactive to physical and emotional stress (Katsu and Iguchi, 2016). More important to the current investigation, however, blood cortisol levels, approximated well by the saliva cortisol level lagging it (Kirschbaum and Hellhammer, 1989), are lowest in the late afternoon up to midnight and begin to rise up again during the second half of the night to peak during the early morning (Pruessner et al., 1997) (Fig. 1a). It has been a long-standing hypothesis that glucocorticoids and their circadian dynamics are linked to cognitive function. There is evidence of a cortisol influence on different cognitive phenomena such as attention, memory formation (Lezak, 1995), and executive functions in general (Roberts et al., 1998).

**Figure 1.**
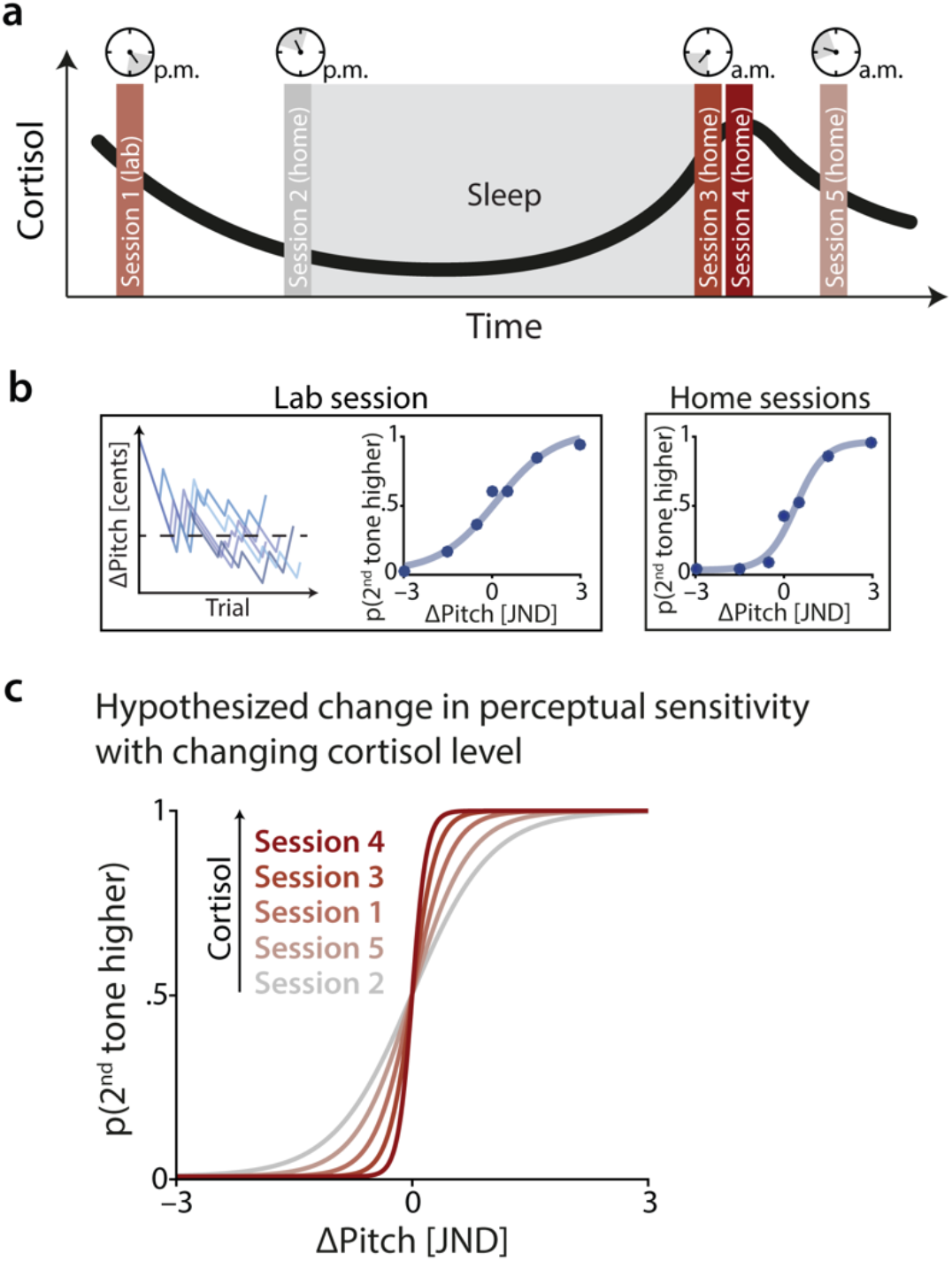
Experimental design and hypothesis. **(a)** Design. In five sessions, participants were asked to take saliva samples, from which their cortisol levels were measured. After a first laboratory session (in the afternoon), participants were asked to perform the other four sessions at home. To capture circadian differences in cortisol levels (black curve), these ‘home’ sessions were timed to align with the individual participant’s sleep-wake cycle such that sessions 2 and 3 had to be completed immediately before going to sleep and immediately after wake-up, respectively. Two further sessions (4 and 5) were performed 30 and 120 minutes after wake-up. **(b)**.Psychophysical testing. In addition to the collection of saliva samples, participants performed a pitch discrimination task in each session. In the lab session, we first assessed individual participants’ pitch discrimination thresholds (just-noticeable difference; JND) using five separate staircases (see Methods for details). These individual JNDs were then used in an online experiment, which participants performed in all five sessions. Psychometric functions (shown in blue) were fit to the data obtained in each session. The slope of the psychometric function served as a measure of perceptual sensitivity. **(c)** Hypothesis. Increased levels of GC availability should result in steeper psychometric functions, reflecting higher perceptual sensitivity. Note that here, sessions are not ordered chronologically but by cortisol level. All illustrations in (a) – (c) are schematic, visualising the hypothesised results of the current study.

Understanding better this potential association between central levels of glucocorticoid hormones and sensory–cognitive performance has implications for the notorious relation of stress-related and hearing disorders (Canlon et al., 2013). It can also further our understanding of how healthy variations in the central availability of stress hormones like cortisol might help regulate sensory and cognitive function.

Generally, we do not know much about how strongly, and in which direction, glucocorticoid availability impacts cognitive function due to a large dynamic range and a variety of pathways by which GCs can act upon central nervous processes (Di et al., 2009; Hall et al., 2015; Lupien and Lepage, 2001; Wolf et al.). Previous results have been mixed: for instance, high baseline cortisol levels have been associated with impaired memory, executive functions, and visual perception (Echouffo-Tcheugui et al., 2018), but also with improved attention and sensory performance in dichotic hearing (Al’Absi et al., 2002). Dijckmans and colleagues (Dijckmans et al., 2017) reported better performance in high cognitive function tasks for participants exhibiting larger variation of cortisol levels throughout the day. An earlier peak and greater magnitude of the typical cortisol awakening response (CAR, a cortisol peak 30–45 min post wake-up) has been shown to be predictive of relatively better executive function-related performance (Evans et al., 2012; Evans et al., 2011).

More generally, it is assumed that a decrease in the dynamic range of circadian GC secretion, either due to an attenuated CAR or due to a slowed elimination of stress-induced cortisol spikes are associated with cognitive impairment in elderly subjects (Beluche et al., 2010; Evans et al., 2011; Stawski et al., 2011). Importantly, little is known on how and to what extent circadian changes in GC availability can influence perceptual processes directly. Visual sensitivity has been reported to fluctuate with time of day (Echouffo-Tcheugui et al., 2018; Stolz et al., 1987; Tassi and Pins, 2009). Clinically, as part of their seminal studies on individuals presenting with GC insufficiency (e.g. characteristic of Addison’s disease) Henkin and colleagues observed a systematic pattern of lowered perceptual thresholds (i.e. better detection) paired with lowered discrimination thresholds (Henkin, 1970), and in the case of audition, a generally lowered dynamic range (Henkin and Daly, 1968). Not least, the systemic administration of synthetic corticosteroids has become a mainstay in treating various hearing disorders, assuming a protective effect of GCs in the inner ear (Trune and Canlon, 2012).

Most directly pertaining to the present study, there is little evidence on how physiological endocrine fluctuations along the circadian cycle influence perception. First evidence with respect to a possible involvement of the circadian system in auditory function is given by the existence of a molecular circadian clock in the cochlea (Meltser et al., 2014) as well as in the inferior colliculus (Park et al., 2016). In addition, Meltser and colleagues (Meltser et al., 2014) reported higher auditory sensitivity, both on molecular and behavioural levels, at specific times of the day (for review see Basinou et al., 2017).

Note that a direct impact of cortisol on auditory perception is physiologically plausible: first, experimental cortisol exposure stimulates the auditory system, but leads to damages in the long term (Al-Mana et al., 2008). Second, GC receptors are expressed in the inner ear, especially in the cochlea (Rarey and Curtis, 1996), as well as in brainstem nuclei involved in auditory processing (Jennes and Langub, 2000). Thus, the SCN-controlled fluctuations in GC availability can impact auditory function at the sensory level directly (Cederroth et al., 2019), and at different levels of the auditory pathway (Canlon et al., 2007).

In the current study, we focus on the impact of the circadian variation of GC availability on auditory perceptual sensitivity in discriminating (i.e., not in detecting the presence of) sounds. We used a psychophysical method, a two-alternative forced-choice (2AFC) task, to describe individual sensitivity for pitch discrimination. Our lead hypothesis here is that GC levels (as proxied by saliva cortisol) impact perceptual performance above and beyond expected drivers such as sex or chronological age: higher levels of saliva cortisol just prior to performing a challenging pitch-discrimination task should lead to a steeper psychometric curve; indicating a state of elevated perceptual sensitivity and, thus, better auditory discrimination abilities. As auxiliary hypotheses, we expected older participants (i) to show less perceptual sensitivity in auditory pitch-discrimination (e.g., Clinard et al., 2010) and (ii) to present with lower levels of saliva cortisol (Evans et al., 2011). The current design allowed us to control for potential confounds of cross-sectional age differences when studying GCs and auditory perception.

We tested a large, age-varying sample of participants to investigate the relationship of saliva cortisol and perceptual performance at the state (i.e., within individuals) and trait level (i.e., between individuals). In detail, we tested a cohort of healthy young adults and a cohort of middle-aged to older participants at five different measurements covering a time interval of approximately 18 hours (see Fig. 1a). We recorded individual sleep duration, and aligned cortisol sampling and behavioural testing relative to the sleep–wake cycle to optimally capture the post-awakening rise and subsequent drop in GC levels (Clow et al., 2010; Wilhelm et al., 2007).

## Results

We investigated the impact of circadian variation in cortisol levels on perceptual sensitivity in a challenging auditory pitch discrimination task. Task difficulty was titrated based on the individual just-noticeable difference (JND). We used separate linear mixed-effects models (i) to test how salivary cortisol secretion changes as a function of time relative to the sleep-wake cycle and age cohort, and (ii) to understand how the observed fluctuation in cortisol levels, in turn, predict perceptual discrimination sensitivity, represented by the slope of the psychometric function. Each model also tested for effects of additional, potential confounders such as sex or sleep duration.

### Explaining momentary states of saliva cortisol

As revealed by model comparison, the momentary level of salivary cortisol was well accounted for by the daytime of measurement (expressed relative to the individual wake-up time and modelled using polynomial regressors of 1^st^, 2^nd^, and 3^rd^ order) and total sleep duration between measurements (conditional R^2^=.75; see Table S1 for full model details). Increased sleep duration was associated with overall lower levels of cortisol (*β* = –.16, *standard error (SE)* = .04, p = .001, *log Bayes factor_10_ (logBF)* = 3.2) while changes in cortisol over time were best described by a cubic trend (*β* = –.15, *SE* = .07, *p* = .025, *logBF* = –.13). As shown in Fig. 2a, this cubic trend captures the decline in cortisol levels from afternoon to late evening and the characteristic cortisol awakening response (CAR; see Fig. 2c). The considerable improvement of model fit by the inclusion of session-specific random intercepts further attests to the impact of daytime on cortisol level (likelihood ratio test; *χ*^*2*^*1* = 69.5, *p* < .001, *logBF* = 31.9). Overall levels of cortisol did not differ significantly between the younger and older cohort (*χ*^*2*^*1* = 2.05, *p* = .15, *logBF* = –1.9; see Fig. 2b). Neither did the cortisol awakening response exhibit a clear effect of age-cohort (Fig. 2c). The inclusion of participants’ sex did not improve model fit, either (*χ*^*2*^*1* = 1.0, *p* = .31, *logBF* = –2.4).

**Figure 2.**
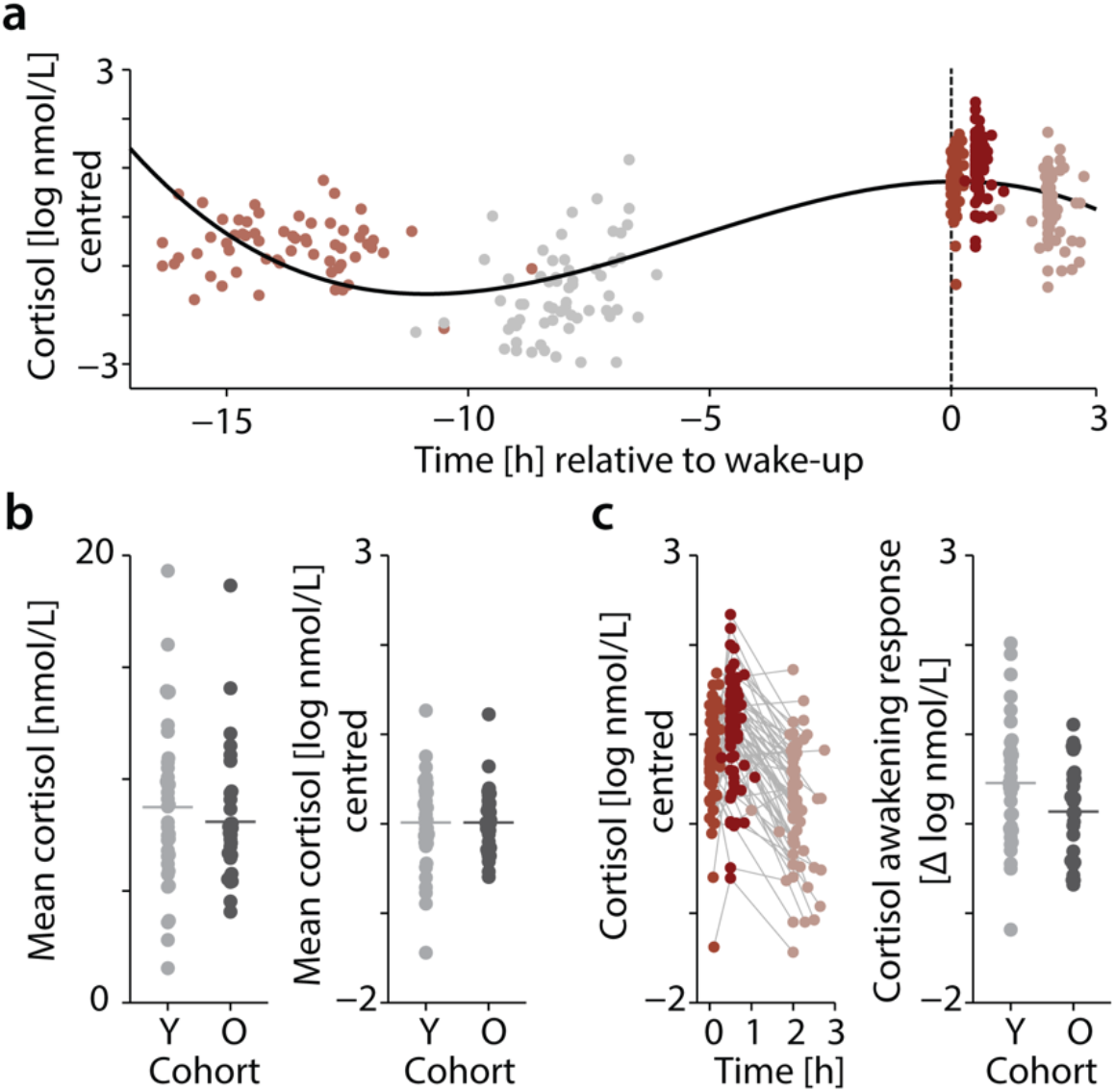
Momentary states of glucocorticoid levels (salivary cortisol) **(a)** Changes in individual salivary cortisol concentration measured in log nmol/L across five experimental sessions. Cortisol levels are mean-centred across all N=68 participants. Sessions are grouped by colour and aligned by wake-up time (dashed vertical line). Black curve shows the cubic trend of time that was modelled using polynomial regression. **(b)** Left panel: individual mean cortisol levels [nmol/L] across sessions shown separately for the younger (Y, light grey) and older (O, dark grey) age cohort. Dots represent individual mean values (N=68), horizontal lines show the respective group average. Right panel: individual mean cortisol levels per group after log-transformation and mean centring for statistical analysis. **(c)** Left panel: trajectory of individual cortisol levels [log nmol/L] following wake-up. Time is expressed relative to wake-up time. Note the rise in cortisol levels 30 min after wake-up (session 4, light teal). Right panel: individual cortisol awakening response (CAR) expressed as the difference in cortisol levels [log nmol/L, centred] 30 min after wake-up relative to wake-up shown separately for the younger (Y, light grey) and older (O, dark grey) age cohort. Horizontal lines indicate the group mean.

### Saliva cortisol predicts perceptual discrimination sensitivity

As the main analysis (Fig. 3), we probed the predictive power of cortisol levels measured just prior to performing a challenging pitch-discrimination task on participants’ perceptual discrimination sensitivity. As indicated by the best-fitting linear mixed-effects model, perceptual sensitivity, operationalized as the slope of the psychometric function, was significantly influenced by the momentary level of cortisol, age cohort and sex (conditional *R*^*2*^ = .47; see Table S2 for full model details).

**Figure 3.**
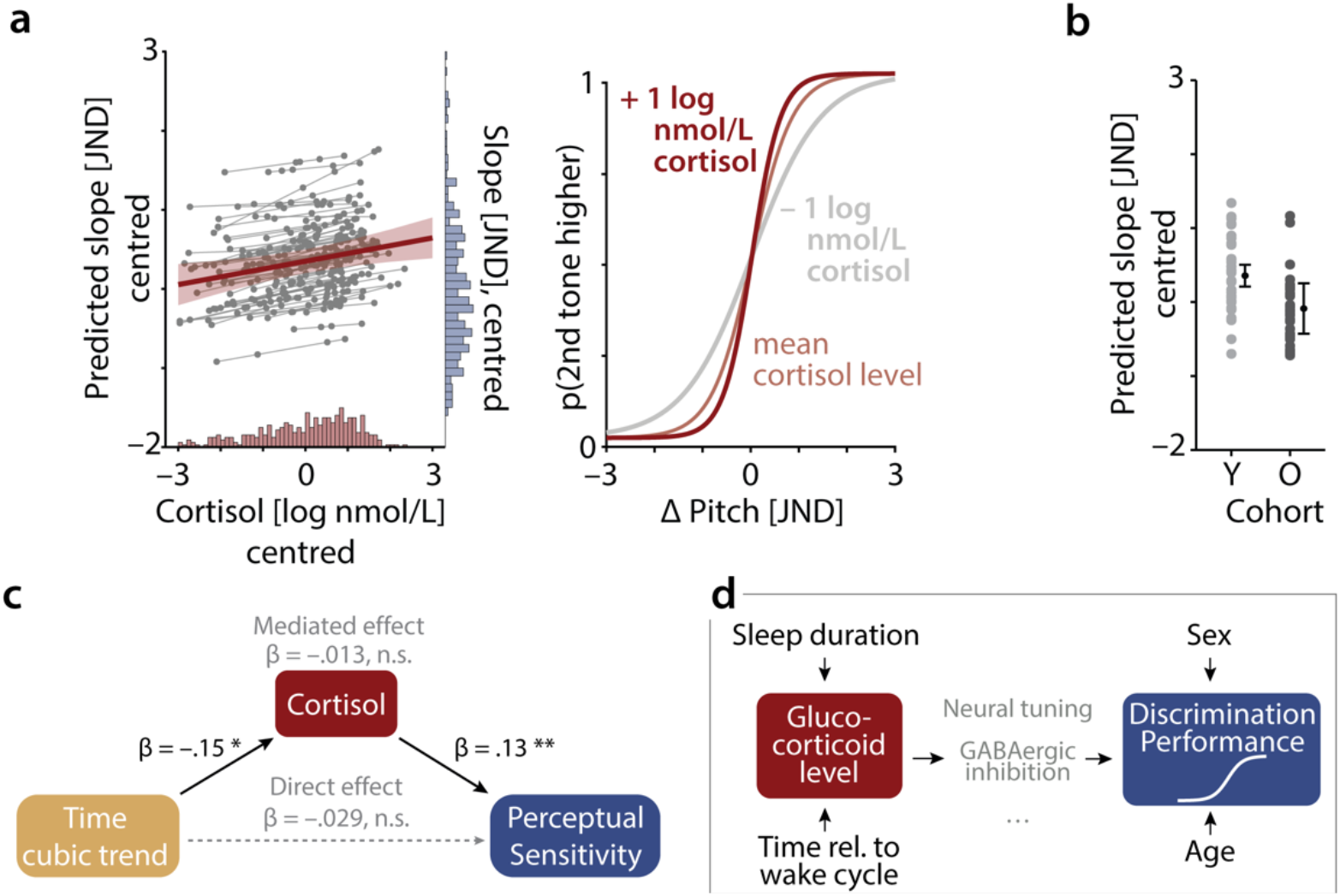
Salivary cortisol predicts perceptual discrimination sensitivity. **(a)** Left panel: change in perceptual sensitivity (operationalised by the slope of psychometric function) as predicted by cortisol. Predicted group-level fixed-effect (red slope) with 95% confidence interval (CI) error band is shown along with the estimated subject-specific random slopes (thin grey lines) and single-subject, single-session predictions (grey dots). Note that subject-specific random slopes did not improve the model fit and were added for illustrative purposes only. Histograms on the bottom and right side of the plot display the distribution of log-transformed cortisol and raw slope values, respectively. Right panel: illustration of how variation in cortisol level impacts the steepness of the psychometric curve. **(b)** Difference in perceptual sensitivity between age groups. Coloured dots (light grey, young (Y) cohort; dark grey, older (O) cohort) show single-subject predicted slope values based on the best-fitting linear mixed-effects model. Black dots represent the fixed-effect group-level prediction and 95% CI. **(c)** Results of causal mediation analysis. Formally accounting for the potentially mediating role of cortisol does not lead to a significant change in the effect of the cubic trend of time on perceptual sensitivity. **(d)** Summary of effects observed. The panel summarises observed (black solid) and statistically excluded (absence of arrows) effects. Intervening (i.e., mediating) effects of *how* GCs can act upon resulting perceptual outcomes must obviously exist, but remain subject to future experimentation. For illustration only, viable paths via a sharpening of neural tuning and/or increased levels of GABAergic inhibition are shown in grey.

In line with our hypotheses, increased levels of cortisol led to heightened perceptual sensitivity (*β* = .13, *SE* = .04, *p* = .004, *logBF* = 1.4). More specifically, as illustrated in Figure 3a (right panel), an increase in cortisol by one unit log(nmol/L) steepened the slope of the psychometric curve by one tenth of the just-noticeable difference (JND). To interpret this effect with respect to the measured log cortisol levels, we examined the range of cortisol levels recorded per individual. On average, the individual cortisol levels changed by 2.71 nmol/L (± sd 0.85 nmol/L) across the course of the experiment. Importantly, as shown by subject-specific slopes added for illustrative purposes in Figure 3a (left panel), the relationship of cortisol and perceptual sensitivity was consistently observable across individual participants. An additional analysis including separate regressors for the state-(i.e., within-subjects) and trait-level (i.e., between-subjects) effect of cortisol on perceptual sensitivity provided additional support for cortisol-driven changes in perceptual sensitivity at the level of the individual participant (within-subject effect of cortisol: *β* = .13, *SE* = .04, *p* = .004, *logBF* = 1.2; between-subject effect cortisol: *β* = .13, *SE* = .23, *p* = .56, *logBF* = –2.7; see Methods and Table S3 for full model details).

As expected, we observed a significant decrease in perceptual sensitivity for the older compared to the younger cohort (*β* = –.52, *SE* = .18, *p* = .005, *logBF* = 1.3; see Fig 3b). More precisely, we observed shallower slopes for older participants with an overall difference in the slopes of younger and older participant of nearly half a JND. Participants’ sex proved to be an additional significant predictor with females showing overall lower perceptual sensitivity (*β* = –.36, *SE* = .18, *p* = .049, *logBF* =–.84; see Figure S1). Participants’ sleepiness [assessed via the Karolinska Sleepiness Scale; see Methods] or response bias [indicated by the point of subjective equality (PSE) on the psychometric function], however, did not influence behavioural performance. The inclusion of these predictors did not significantly improve model fit (likelihood ratio tests, all *p* > .067, all *logBFs* < –1.2).

Lastly, we investigated whether changes in cortisol would differentially impact perceptual sensitivity across the two age groups, despite overall comparable levels of cortisol observed for younger and older adults. However, the inclusion of the respective interaction term did not improve the model fit (*χ*^*2*^*1* = .91, *p* = .34, *logBF* = –2.4).

### Cortisol does not impact response bias

To investigate whether the impact of momentary cortisol levels was specific to perceptual sensitivity, we ran a control model probing for their effect on response bias. We found a significant increase in PSE for older participants (*β* = .44, *SE* = .2, *p* = .027, *logBF* = –.34). Importantly, however, circadian fluctuations in cortisol did not significantly predict changes in response bias (*β* = .001, *SE* = .04, *p* = .98, *logBF* = –2.9, see supplemental Fig. S3 and Table S4).

### Ruling out confounding effects of task proficiency

One concern we aimed to target is the obvious repetition of the pitch discrimination task in close succession, especially in the morning of the second testing day (i.e., three times of testing within approximately two hours). Reassuringly, however, no training or time-of-day effects on our main outcome measure of pitch-discrimination perceptual sensitivity were evident (Fig. S2) as the inclusion of session (χ^2^1 = 1.6, p = .21, logBF = –2.1) or time (linear, quadratic, cubic trend) did not improve model fit (all p > .4, logBFs < –2.5).

### Cortisol is not simply mediating an effect of daytime on perceptual sensitivity

An additional control analysis considered the possibility that the observed link between cortisol and perceptual discrimination sensitivity could reflect an indirect effect of daytime on perceptual sensitivity. While the absence of any systematic changes in the slope of the psychometric function with time (see above) rendered this scenario unlikely, we still formally tested this possibility using causal mediation analysis. As shown in Figure 3c, the comparison of the estimated total and direct effect of time (cubic trend) on perceptual sensitivity showed a comparably small and non-significant change (–.042, CI[–.13, .04] vs. –.029, CI[– .13, .07]) when accounting for the indirect influence via cortisol (–.013, CI[–.07, .04]). In other words, the observed increase in perceptual sensitivity with increasing levels of salivary cortisol does provide evidence for their potentially causal relationship.

## Discussion

Does the momentary availability of GCs predict changes in perceptual abilities, and if so, to what degree? The current study set out to gather decisive data on this seemingly simple question. In a mixed between-and within-participants design using multiple saliva cortisol samples and multiple associated behavioural assessments of perceptual sensitivity throughout the circadian cycle, we here have indeed provided evidence for a link between GC level and auditory perceptual discrimination ability.

A first result lending overall credibility to our approach is the circadian modulation of saliva cortisol. A highly consistent pattern of relative cortisol level displacement dependent on daytime of measurement was observable (Fig. 2a), which in concert with the individual duration of sleep (taking place between measurements 2 and 3) could explain the observed GC variance to a large degree.

Second, as the main result of our study, saliva cortisol levels just prior to performing the pitch discrimination task were predictive of perceptual sensitivity. Statistically dissecting the influence of trait-level (i.e., person to person) versus state-level (i.e., session to session) variation in cortisol showed that it was the momentary cortisol level just prior to behavioural testing that covaried with perceptual sensitivity. The robustness and size of this effect is illustrated in Figure 3a: a change of one’s own cortisol level by one unit log(nmol/L) steepens one’s psychometric curve by approximately 1/10 of the just-noticeable difference in pitch. Essentially all participants showed this positive relationship of momentary saliva cortisol levels with the steepness of the psychometric curve in pitch discrimination. Lastly, a series of control analyses underscores the directness and putative causality of the effect of cortisol on auditory discrimination performance.

This result fills various gaps in our knowledge on how the endocrine system impacts perception and behaviour. We will discuss potential mechanisms, limitations, and implications below.

### Potential mechanisms: how could GC levels act upon perceptual sensitivity?

The present results imply that, within normal levels of a healthy endocrine system, relative increases in centrally available GCs are accompanied by an objective improvement in the ability to discriminate sounds. This is broadly in line with a view of stress hormones and activity of the hypothalamic-pituitary-adrenal (HPA) axis as preparing the body for action (Habib et al., 2001). Enhanced discrimination abilities certainly fit into this view.

What are mechanistic pathways by which glucocorticoids could bring such an improved discrimination about, and how specific to auditory discrimination might these pathways be? We here aim to briefly cover two mechanistically conceivable paths – one relating to the established circadian dependence of GC and GC receptor dynamics in the inner ear and auditory system, and one relating to improved neural tuning via potentially GC-related GABAergic signalling.

First, there is ample evidence by now for a causal role of glucocorticoid dynamics in the healthy as well as stress-related malfunction of the auditory system (e.g., Canlon et al., 2007; Canlon et al., 2013; Hossain et al., 2008). The immunosuppressive and anti-inflammatory effect of GCs require binding to glucocorticoid receptors (GRs), which are expressed not only in the structures of the HPA axis but also in the auditory system (Canlon et al., 2007). Also, the fine-tuned balance of GC receptors with mineralocorticoid receptor activity is well-documented (e.g., de Kloet et al., 2018; Singer et al., 2018). Thus, the glucocorticoid dynamics in the inner ear are under circadian control, which has also been demonstrated for the auditory system in the rodent (Cederroth et al., 2019). Accordingly, protective effects of GCs against hearing damage have been proposed. The effects of GC availability on perceptual discrimination abilities observed here might in part reflect such slow-acting variations in peripheral or central auditory function (Cederroth et al., 2019).

Second, GCs have been shown to facilitate inhibitory GABAergic synaptic input (and to concomitantly suppress excitatory glutamatergic drive) at least in hypothalamic neurons, part of the HPA axis (Di et al., 2009). It remains speculative at this point whether such a combined effect of GCs might also tip other brain areas towards inhibition, with concomitant improvements of discrimination abilities both at the neural and behavioural level. This poses a testable pathway: recent neurophysiology work using optogenetic stimulation of layer-6 cortical neurons in rodents has provided compelling evidence for a dissociation and rapid switching of detection-optimal versus discrimination-optimal configurations at neural and behavioural levels (Guo et al., 2017; Linden, 2017).

Both peripheral effects of GC and central effects at various stages need to be considered and tested in detail in future studies. However, a two-alternative forced-choice (2AFC) discrimination task such as the present one requires the system to detect equally well two stimuli and to arbitrate between the two with respect to one task-relevant dimension (here, tone frequency). In signal detection theoretic terms, one stimulus (here, the one higher in tone frequency) is considered as “signal plus some noise” and should be chosen by the listener, while the other is considered “only noise”. Thus, improving sensitivity in such a task requires a mechanism that is able to improve the “signal to noise” ratio— either at the level of neural encoding (inner ear, midbrain, auditory cortex), or at the level of decision-making (auditory cortex and beyond), or both.

A concept viable at all these levels is neural tuning, the degree to which a neuron or neuronal population is selectively responsive to a certain range along a given featural dimension, here, pitch or sound frequency. Neural tuning in auditory cortex is known to be highly adaptive to task demands in any given listening situation (Atiani et al., 2009; Holdgraf et al., 2016; Jiang et al., 2018). Additionally, improved discriminability of tones is a phenomenon with a clear auditory-cortical contribution (Christensen et al., 2019). Recent work in humans also underscores that ongoing neural population dynamics, which should be especially amenable to endocrine modulation, can flexibly (i.e., from trial to trial) affect behavioural sensitivity (e.g., Gelbard-Sagiv et al., 2018; Waschke et al., 2019).

It is worth noting that an inhibition-and tuning-related mechanism has received at least circumstantial support by the seminal clinical observations of Henkin and colleagues in the late 1960s. Primary GC insufficiency as observed in Addison’s disease was found accompanied by paradoxically improved detection thresholds but a decrease in discrimination abilities (Henkin and Daly, 1968; Henkin et al., 1967). There thus lies great promise in better understanding the differential GC susceptibility of these detection and discrimination processes in the auditory system.

In sum, an obvious next step should be to manipulate GC availability in the human listener directly. In healthy participants, a relatively unspecific but carefully titrated administration of synthetic GC analogues can easily be used to obtain experimental control over GC levels within the normal dynamic range of HPA axis activity (Born et al., 1987). Not least, this would bring the field back to a promising lead that it left seemingly behind half a century ago: Namely, the seminal work by Henkin and others on mechanistic pathways how adrenocortical hormones control the detection and integration of sensory signals (Henkin, 1970). In clinical patients with primary GC insufficiencies (e.g. Addison’s disease) or GC hyper-availability (Cushing’s disease), it will be fruitful to build on pioneering but technically limited work linking lowered thresholds (e.g., hyper-sensibility in detection) but lowered discrimination abilities in the auditory domain to cortisol states (e.g., Henkin and Daly, 1968).

### Potential confounders of a causal influence of cortisol on perceptual sensitivity

As summarised in Figure 3d, the current work helps us rule out two potential (i.e., theoretically plausible) confounders. Namely, both participants’ age and participants’ sex were indeed predictive of perceptual performance (with younger and male participants outperforming older and female participants, respectively). However, they are both highly unlikely to confound the observed effect of GCs on this performance, as neither of the two could account for momentary cortisol levels (note the absence of arrows from sex or age into GC in Fig. 3d).

Not shown in Figure 3d, but reported in detail above, other more global “state” variables such as time of day (recall that most data here were acquired either just prior to bedtime or immediately after waking up) or total duration of sleep were good predictors of the momentary cortisol level. These, however, failed to account for any meaningful variance in the behavioural outcome (note the absence of arrows from *Time relative to wake cycle* and *Sleep duration* to *Discrimination performance*). Thus, it was not the case that, for example, participants who had slept more were overall providing higher perceptual sensitivity across all testing instances or vice versa. Neither did testing in the evening yield lower discrimination performance, all other things being equal.

Unsurprisingly, the mediation analysis we performed for the sake of completeness (see Results) also did not provide any evidence for a potential mediation (i.e., daytime → cortisol → performance). Instead, our results expose a more direct link from momentary salivary cortisol level to sensitivity in perceptual discrimination.

Note that we made a set of design choices (e.g., cortisol sampling always directly preceding the behavioural test) that help to rule out a conceivable, reverse causal relationship (i.e., worse performance in the behavioural task leading to perceived stress and, thus, to higher cortisol level). Such a hypothesis, however, is rendered unlikely on two grounds. First, a previous study has found no effect of task effort or hearing status on cortisol as a marker of stress (Zekveld et al., 2019). Second, the present data themselves invalidate this notion, as higher cortisol levels just prior to testing were accompanied by better, not worse performance at test.

This leaves us with one potential unobserved confound, namely, arousal. Could elevated levels of arousal have led to higher cortisol levels and, hence, to better behavioural performance? Arousal is generally assumed to establish an inverted u-shaped impact on performance (i.e., the “Yerkes-Dodson law”; Gelbard-Sagiv et al., 2018; Waschke et al., 2019; Yerkes and Dodson, 1908). However, the fact that we did not observe such a pattern does not necessarily rule out a confounding influence of arousal as our paradigm might have captured only activity along the “rising” flank of such an inverted u.

Nevertheless, we deem a confounding influence of autonomic arousal on both, GC levels and performance, unlikely on various grounds. First, the relationship of stress on individual cortisol levels is not as clear-cut as often assumed (Kirschbaum et al., 1995). Second, autonomic arousal markers, such as heart rate and blood pressure on the one hand and cortisol on the other, have been shown to exert a dissociable, that is, sufficiently independent impact on performance in a verbal memory task (Schwabe et al., 2008). Third, even if we assume a mechanistic link between arousal and cortisol, it is hard to imagine why autonomic arousal should have covaried so consistently across our diverse cohorts of young and old adults and time of day in order to yield the consistent behavioural effects. The Karolinska Sleepiness Scale, assessed here as a control measure and arguably a proxy of arousal did not indicate any systematic covariation with behavioural outcome.

Nevertheless, a future study co-registering pupil dilation (as an established proxy of arousal) or physiological markers such as skin conductance and heart rate, ideally in a setting where GC levels are manipulated experimentally, should help to illuminate the causal links between arousal, cortisol, and discrimination performance.

### Implications

We here have shown that a main neurobiological circadian rhythm in the human body, the secretion of glucocorticoids (here, captured as saliva cortisol), covaries with the individual fluctuation of auditory perceptual abilities (here, captured as pitch-discrimination sensitivity immediately after taking the saliva sample). We have demonstrated that momentary GC levels show the expected circadian change, and that these within-individual fluctuation of GC levels directly predict perceptual discrimination sensitivity.

This result opens at least two new research avenues: first, experimental control and manipulation of endocrine modulators such as the GC system can help to constrain future research into the organisation of the auditory system. Second, our study opens new paths to improving or restoring discrimination abilities, a particularly vulnerable aspect of auditory function in both ageing generally and age-related hearing-loss specifically.

## Materials and methods

### Participants

Seventy-five participants took part in this study, acquired in two waves (younger participants in April–May 2018, older participants in April–May 2019). The participants were recruited through the database of the Department of Psychology at the University of Lübeck, using the online recruiting system ORSEE (Greiner, 2015). The cohort of younger participants consisted of 37 university students (24 females, mean age 22.6, SD = 2.58, age range 19–30 years). The cohort of middle-aged and older participants consisted of 38 persons (19 females, mean age 60.6, SD = 5.98, age range 50–70 years); 16 of them were retired.

All participants were screened to avoid any history of disorders that could have impacted their GC balance, such as neurological or psychiatric disorders as well as any known metabolic diseases. Furthermore, none showed a BMI over 30 kg/m^2^ or had been working in shifts. None reported any known hearing disorders, severe current hearing loss, or a persistent tinnitus. Note, however, that participants with mild age-related hearing loss were not excluded from the cohort of older participants due to its high prevalence in this age group.

In the cohort of younger participants, none took any medication that could have influenced their GC balance, including medication for asthma-or allergy treatment, systemic immunosuppressants or antihypertensives.

In the cohort of older participants, more lenient inclusion criteria with respect to medication applied (see Table S5). Here, participants who took any type of antihypertensives were still included to allow for a representative sample of older adults.

Written informed consent was collected from all participants according to procedures approved by the Research Ethics Committee of the University of Lübeck. Listeners were paid 25–30 € or received course credit for their participation in the experiment.

### Experimental protocol

On the first day, participants came to the laboratory between 4pm and 6pm for the first session, lasting about one and a half hours. A maximum of four participants conducted the first session on a given day. The session started with an adaptive tracking procedure that measured auditory pitch thresholds (see section Psychoacoustic testing for details). Participants were then asked to complete three questionnaires on their general medical history, their chronotype (Horne and Ostberg, 1976), and their momentary sleepiness (assessed using the Karolinska Sleepiness Scale; Akerstedt and Gillberg, 1990). The scale consists of three items: (1) sleepiness during the last 10 minutes (nine steps on a Likert-Scale), (2) the current state with relaxation on one end and tension on the other end of a visual analogue scale and (3) the current fatigue (visual analogue scale).

Next, participants received detailed instructions for the subsequent measurements. Each session included taking a saliva sample and performing a challenging pitch discrimination task in a browser-based online study (Labvanced, Osnabrück), followed by the sleepiness questionnaire. According to their auditory pitch threshold, participants were assigned to an experimental group, designed to yield equivalent difficulties of the pitch discrimination task (see *Assessment of pitch discrimination thresholds* below), and provided with an individual link, which gave them exactly five times access to the online task.

Finally, participants completed the first session: taking a saliva sample first (see Saliva cortisol collection for details) and performing the online pitch discrimination task secondly before they were sent home. Throughout all sessions, participants in the younger cohort used their own technical devices (laptop and headphones) whereas participants in the older cohort used their own headphones for all experimental sessions but were provided with computers for the first session due to their lack of portable computers. Usage of participants’ own equipment ensured that the acoustic properties of the pitch discrimination task remained constant across sessions and, whenever possible, that the experiment could be adequately performed with the participants’ personal equipment.

All other measurements were conducted at home, scheduled at certain times of day relative to the participants’ sleep–wake cycle: Session 2 had to be performed just before going to sleep, Session 3 immediately after waking up (participants were instructed to place the equipment, or at least the Salivette tube for the saliva sample, next to their bed), Session 4 30 minutes, and Session 5 about 120 minutes after awakening. To assess compliance and to gather information about the time of events, participants recorded the starting time of each session as well as the activities that they were engaged in between two consecutive sessions in a time protocol. Additionally, they were asked to maintain their typical sleeping and wake-up times, which they had recorded for the last two weeks.

### Saliva cortisol collection

Salivary cortisol level was measured to deduce the amount of unbound cortisol in blood (Kirschbaum and Hellhammer, 1994). To capture a comprehensive cycle of cortisol secretion as used in former studies (Dijckmans et al., 2017; Evans et al., 2011; Lee et al., 2007), including the characteristic morning rise, a saliva sample was collected at each single experimental session. As described above, sessions were scheduled according to the individual participant’s wake-up time. Following instructions and the collection of a first saliva sampling in the lab session, participants were provided with a saliva self-collection pack containing four Salivette Cortisol tubes (Sarstedt, Nümbrecht, Germany), pre-labelled with participant code and number of session, and written instructions. For a correct usage, the Salivette dental swab from the correctly labelled Salivette had to be chewed until fully saturated and then be put back into the tube. Saliva samples were then stored in the participants’ own freezer until they were brought back or picked up after one to seven days, together with the time protocol and stored in the freezer of the Department of Psychology.

To avoid bias, participants were asked not to smoke, eat, drink (except water) or brush their teeth 30 minutes before sampling.

All saliva samples (180 from the younger cohort and 185 from the older cohort) were analysed at the Biochemical Laboratory of the Technical University Dresden. The fraction of free cortisol in saliva (salivary cortisol) was determined using a time-resolved immunoassay with fluorometric detection (for detailed method see Dressendorfer et al., 1992) and reported back to the authors in the unit of measurement, nmol/l, to 1-decimal precision.

### Psychoacoustic testing

#### Assessment of pitch discrimination thresholds

In the first session, we assessed individual participants’ pitch discrimination thresholds (i.e., their so-called just-noticeable differences; JNDs) using a weighted one-up, one-down method (Kaernbach, 1991). On each trial, participants heard two pure tones. Each tone had a duration of 100 ms with a silence period of 25 ms between tones. The first tone always had a frequency of 1 kHz; the frequency of the second tone differed from that of the first tone by delta f. The participants were asked to indicate via button press which of the two tones had the higher frequency. The next trial started 750 ms after the participants’ response. Responses were self-paced. No feedback was given.

The assessment of pitch discrimination thresholds comprised five staircases per participant. Each staircase started with a delta f of 100 cents (i.e., one semitone). In the first phase, delta f was increased by a factor of 2.25 following an incorrect response and was decreased by the cube root of 2.25 following a correct response. Hence, the magnitude of upward steps was three times larger than the magnitude of downward steps, estimating approximately 75%-correct on the psychometric function. In the second phase, we used a factor of 1.5 and cube root of 1.5 for up-and down-steps, respectively. Each staircase was terminated after the twelfth reversal; there were four reversals in the first phase and eight reversals in the second phase. The threshold in each staircase was defined as the arithmetic mean of delta fs visited on all second-phase reversal trials. Finally, individual JNDs were defined as the average of thresholds across all five staircases per participant.

### Assessment of psychometric curves

In each of the five sessions, participants performed a pitch discrimination task in a browser-based online study (Labvanced, Osnabrück). This task was similar to the assessment of pitch discrimination thresholds, which was completed in the first session only (see above): on each trial, participants heard two pure tones which differed in frequency and were asked to indicate which tone had the higher pitch. Here, however, we used a method of constant stimuli to assess participants’ individual pitch sensitivity. In each session, participants completed 148 trials, comprising seven stimulus levels relative to their individual pitch discrimination threshold (JND). This means that participants were assigned to different groups based on their individual thresholds to ensure similar difficulty levels across participants. We considered five different groups: 5ct, 10ct, 15ct, 20ct, and 25ct. Participants were assigned to the group closest to their individual JND (e.g., a participant with a JND of 7.5ct would be assigned to the 10ct group, while a participant with a JND of 7.4ct would be assigned to the 5ct group).

The stimulus levels were approximately -3, -1.5, -0.5, 0, 0.5 1.5, and 3 JNDs. This choice of stimulus levels allowed us to sample the linear part of the logistic function (slope), while also capturing its asymptotes (Herbst and Obleser, 2019; Waschke et al., 2019). Note that a stimulus level of zero JND means that the two tones on a given trial had the same frequency of 1 kHz. Hence, there was no correct response for this stimulus level. Each stimulus level was presented 21 times per session. We additionally included one dummy trial at the beginning of each session. The response in this trial was excluded from the analysis; however, inclusion of this dummy trial allowed us to present the stimulus levels using a type-1 index-1 sequence (Finney and Outhwaite, 1956). Type-1 index-1 sequences control for potential carry-over effects by first-order counterbalancing. This means that each stimulus level has the same probability to occur after each other stimulus level, including itself.

In each session, we calculated the proportion of ‘second tone higher’ responses per stimulus level and fitted a logistic function to the data using the Palamedes toolbox version 1.7.0 (Prins and Kingdom, 2018) in MATLAB (MathWorks, Natick, Massachusetts, USA; R2017b). We fitted three parameters: The point of subjective equality (PSE; i.e., the point where subjects reported ‘second tone higher’ in 50% of trials), the slope at the PSE (i.e., our measure of perceptual sensitivity), and the lapse rate (i.e., the lower asymptote). The guess rate (i.e., the higher asymptote) was fixed at 1 minus the guess rate, which resulted in symmetric asymptotes of the psychometric fit.

Data sets from eight individual sessions did not follow a psychometric curve and no fit was possible. Additionally, we excluded fits with extreme slopes (i.e., larger than 5) as well as flat psychometric curves. Based on these criteria, six participants produced less than two usable fits. All data from these participants were therefore excluded from further analyses.

The data from one participant in the younger cohort who reported to follow an unusually shifted sleep-wake cycle were excluded prior to analysis. The data of six participants in the older cohort were excluded from analysis because they either dropped out of the study after the first session (N=3), or because of missing or unusable data in more than three sessions (N=3; see details on psychoacoustic testing below).

The final sample consisted of N=68 individuals and, in sum, we used 318 of a possible maximum of 340 observations in the statistical analyses.

### Statistical analysis

We used linear mixed-effect models to investigate how circadian fluctuations in salivary cortisol level influence perceptual sensitivity. To this end, we first investigated how cortisol expression levels change throughout the day by modelling increasingly complex trajectories via the inclusion of polynomial regressors of different orders. We also tested for changes in cortisol levels as a function of sleep duration, age cohort (young/old), and sex (male/female). In the main analysis, we then modelled the influence of momentary cortisol levels on auditory perceptual discrimination sensitivity, expressed as the slope of the psychometric function. We also tested for the impact of time (expressed relative to the individual wake-up time), age cohort, sex, sleep duration, pitch group, sleepiness (assessed using the Karolinska Sleepiness Scale), and response bias (expressed as the point of subjective equality on the psychometric curve, PSE).

Estimation and selection of linear mixed-effect models (Gaussian distribution, identity link function) followed an iterative model fitting procedure (Alavash et al., 2019; Tune et al., 2018). We started with an intercept-only null model including subject-specific random intercepts and added fixed-effects terms in a stepwise procedure following their conceptual importance. Main effects were added prior to higher-order interaction terms. Lastly, we tested whether the inclusion of a session-specific random intercept or subject-specific random slopes for time-varying within-subject effects would improve model fit. Change in model fit was assessed via likelihood ratio tests on models (re-fit with maximum-likelihood estimation for comparison of fixed effects).

We used deviation coding for categorical predictors. Single-subject observations with unusually high cortisol levels of above 60 nmol/L were discarded. Cortisol levels were log-transformed and as all other continuous variables z-scored prior to modelling. To facilitate interpretation, in the visual presentation of model results, we transformed the continuous variables back to their original units.

An additional control analysis included two separate predictors for the influence of cortisol on perceptual sensitivity to tease apart within-and between-subject effects of cortisol on behaviour. Mean cortisol levels per subject captured the trait-like, between-subject effect while the state-like, within-subject effect was modelled by the session-by-session deviation from this subject-level mean (Bell et al., 2019).

In a second control analysis, we performed a causal mediation analysis (Imai et al., 2010) to formally test the possibility of cortisol only mediating a daytime effect on perceptual sensitivity. We estimated the direct, indirect (mediated) and total effect of the cubic trend of time on perceptual sensitivity using the same set of covariate regressors in the mediation and outcome model. We calculated 95 % quasi-Bayesian confidence intervals using 5,000 replications.

We report p-values for individual model terms that were derived using the Kenward-Roger approximation for degrees of freedom (Luke, 2017). As goodness-of-fit measures, we report R^2^ (marginal and conditional R^2^; taking into account only fixed or fixed and random effects, respectively) along with the Akaike information criterion (AIC) (Nakagawa et al., 2017). To facilitate interpretation of (non-)significant effects, we also calculated the Bayes factor (BF) based on the comparison of Bayesian information criterion (BIC) values as proposed by Wagenmakers (Wagenmakers, 2007). Throughout we report log Bayes Factors, with a log BF of 0 representing equal evidence for and against the null hypothesis; log BFs with a positive sign indicating relatively more evidence for the alternative hypothesis than the null hypothesis, and vice versa. Magnitudes > 1 are taken as moderate, > 2.3 as strong evidence for either of the alternative or null hypotheses, respectively. All analyses were performed in R (version 3.6.1) using the lme4 (Bates et al., 2015), mediation (Tingley et al., 2014), and sjPlot (Lüdecke, 2020) packages.

### Data availability

All data and scripts for data analysis will be made available at Open Science framework (OSF; https://osf.io/ns26m/) upon publication.

## Acknowledgments

This research has been supported by the European Research Council (ERC-CoG-2014-646696 “AUDADAPT” to JO) and the Deutsche Forschungsgemeinschaft (DFG OS353-10/1 to HO). Franziska Scharata and Anne Hermann helped acquire the data.

## References

Akerstedt, T., and Gillberg, M. (1990). Subjective and objective sleepiness in the active individual. Int J Neurosci 52, 29–37.

Al-Mana, D., Ceranic, B., Djahanbakhch, O., and Luxon, L.M. (2008). Hormones and the auditory system: A review of physiology and pathophysiology. Neuroscience 153, 881–900.

Al’Absi, M., Hugdahl, K., and Lovallo, W.R. (2002). Adrenocortical stress responses and altered working memory performance. Psychophysiology 39, 95– 99.

Alavash, M., Tune, S., and Obleser, J. (2019). Modular reconfiguration of an auditory control brain network supports adaptive listening behavior. Proc Natl Acad Sci U S A 116, 660–669.

Atiani, S., Elhilali, M., David, S.V., Fritz, J.B., and Shamma, S.A. (2009). Task difficulty and performance induce diverse adaptive patterns in gain and shape of primary auditory cortical receptive fields. Neuron 61, 467–480.

Basinou, V., Park, J.S., Cederroth, C.R., and Canlon, B. (2017). Circadian regulation of auditory function. Hear Res 347, 47–55.

Bates, D., Mächler, M., Bolker, B., and Walker, S. (2015). Fitting Linear Mixed-Effects Models Using lme4. Journal of Statistical Software 67, 1–48.

Bell, A., Fairbrother, M., and Jones, K. (2019). Fixed and random effects models: making an informed choice. Quality \& Quantity 53, 1051--1074.

Beluche, I., Carrière, I., Ritchie, K., and Ancelin, M.L. (2010). A prospective study of diurnal cortisol and cognitive function in community-dwelling elderly people. Psychological Medicine 40, 1039–1049.

Born, J., Kern, W., Fehm-Wolfsdorf, G., and Fehm, H.L. (1987). Cortisol effects on attentional processes in man as indicated by event-related potentials. Psychophysiology 24, 286–292.

Canlon, B., Meltser, I., Johansson, P., and Tahera, Y. (2007). Glucocorticoid receptors modulate auditory sensitivity to acoustic trauma. Hear Res 226, 61–69.

Canlon, B., Theorell, T., and Hasson, D. (2013). Associations between stress and hearing problems in humans. Hear Res 295, 9–15.

Cederroth, C.R., Park, J.S., Basinou, V., Weger, B.D., Tserga, E., Sarlus, H., Magnusson, A.K., Kadri, N., Gachon, F., and Canlon, B. (2019). Circadian Regulation of Cochlear Sensitivity to Noise by Circulating Glucocorticoids. Curr Biol 29, 2477–2487 e2476.

Christensen, R.K., Linden, H., Nakamura, M., and Barkat, T.R. (2019). White Noise Background Improves Tone Discrimination by Suppressing Cortical Tuning Curves. Cell Rep 29, 2041–2053 e2044.

Clinard, C.G., Tremblay, K.L., and Krishnan, A.R. (2010). Aging alters the perception and physiological representation of frequency: evidence from human frequency-following response recordings. Hear Res 264, 48–55.

Clow, A., Hucklebridge, F., Stalder, T., Evans, P., and Thorn, L. (2010). The cortisol awakening response: more than a measure of HPA axis function. Neurosci Biobehav Rev 35, 97–103.

de Kloet, E.R., Meijer, O.C., de Nicola, A.F., de Rijk, R.H., and Joels, M. (2018). Importance of the brain corticosteroid receptor balance in metaplasticity, cognitive performance and neuro-inflammation. Front Neuroendocrinol 49, 124–145.

Di, S., Maxson, M.M., Franco, A., and Tasker, J.G. (2009). Glucocorticoids Regulate Glutamate and GABA Synapse-Specific Retrograde Transmission via Divergent Nongenomic Signaling Pathways. The Journal of Neuroscience 29, 393--401.

Dibner, C., Schibler, U., and Albrecht, U. (2010). The Mammalian Circadian Timing System: Organization and Coordination of Central and Peripheral Clocks. Annu Rev Physiol 72, 517–549.

Dijckmans, B., Tortosa-Martínez, J., Caus, N., González-Caballero, G., Martínez-Pelegrin, B., Manchado-Lopez, C., and Clow, A. (2017). Does the diurnal cycle of cortisol explain the relationship between physical performance and cognitive function in older adults? European Review of Aging and Physical Activity 14, 6.

Dressendorfer, R.A., Kirschbaum, C., Rohde, W., Stahl, F., and Strasburger, C.J. (1992). Synthesis of a cortisol-biotin conjugate and evaluation as a tracer in an immunoassay for salivary cortisol measurement. J Steroid Biochem Mol Biol 43, 683–692.

Echouffo-Tcheugui, J.B., Conner, S.C., Himali, J.J., Maillard, P., DeCarli, C.S., Beiser, A.S., and Seshadri, S. (2018). Circulating cortisol and cognitive and structural brain measures: The Framingham Heart Study. Neurology 91, 1961– 1970.

Evans, P., Hucklebridge, F., Loveday, C., and Clow, A. (2012). The cortisol awakening response is related to executive function in older age. International Journal of Psychophysiology 84, 201–204.

Evans, P.D., Fredhoi, C., Loveday, C., Hucklebridge, F., Aitchison, E., Forte, D., and Clow, A. (2011). The diurnal cortisol cycle and cognitive performance in the healthy old. International Journal of Psychophysiology 79, 371–377.

Finney, D.J., and Outhwaite, A.D. (1956). Serially balanced sequences in bioassay. Proc R Soc Lond B Biol Sci 145, 493–507.

Gelbard-Sagiv, H., Magidov, E., Sharon, H., Hendler, T., and Nir, Y. (2018). Noradrenaline Modulates Visual Perception and Late Visually Evoked Activity. Curr Biol 28, 2239–2249 e2236.

Greiner, B. (2015). Subject Pool Recruitment Procedures: Organizing Experiments with ORSEE. Journal of the Economic Science Association 1, 114--125.

Guo, W., Clause, A.R., Barth-Maron, A., and Polley, D.B. (2017). A Corticothalamic Circuit for Dynamic Switching between Feature Detection and Discrimination. Neuron 95, 180–194 e185.

Habib, K.E., Gold, P.W., and Chrousos, G.P. (2001). Neuroendocrinology of stress. Endocrinol Metab Clin North Am 30, 695-728; vii-viii.

Hall, B.S., Moda, R.N., and Liston, C. (2015). Glucocorticoid Mechanisms of Functional Connectivity Changes in Stress-Related Neuropsychiatric Disorders. Neurobiol Stress 1, 174–183.

Henkin, R.I. (1970). The effects of corticosteroids and ACTH on sensory systems. Prog Brain Res 32, 270–294.

Henkin, R.I., and Daly, R.L. (1968). Auditory detection and perception in normal man and in patients with adrenal cortical insufficiency: effect of adrenal cortical steroids. J Clin Invest 47, 1269–1280.

Henkin, R.I., McGlone, R.E., Daly, R., and Bartter, F.C. (1967). Studies on auditory thresholds in normal man and in patients with adrenal cortical insufficiency: the role of adrenal cortical steroids. J Clin Invest 46, 429–435.

Henry, M.J., and Obleser, J. (2012). Frequency modulation entrains slow neural oscillations and optimizes human listening behavior. Proc Natl Acad Sci U S A 109, 20095–20100.

Herbst, S.K., and Obleser, J. (2019). Implicit temporal predictability enhances pitch discrimination sensitivity and biases the phase of delta oscillations in auditory cortex. NeuroImage 203, 116198.

Holdgraf, C.R., de Heer, W., Pasley, B., Rieger, J., Crone, N., Lin, J.J., Knight, R.T., and Theunissen, F.E. (2016). Rapid tuning shifts in human auditory cortex enhance speech intelligibility. Nat Commun 7, 13654.

Horne, J.A., and Ostberg, O. (1976). A self-assessment questionnaire to determine morningness-eveningness in human circadian rhythms. International Journal of Chronobiology 4, 97–110.

Hossain, A., Hajman, K., Charitidi, K., Erhardt, S., Zimmermann, U., Knipper, M., and Canlon, B. (2008). Prenatal dexamethasone impairs behavior and the activation of the BDNF exon IV promoter in the paraventricular nucleus in adult offspring. Endocrinology 149, 6356–6365.

Imai, K., Keele, L., and Tingley, D. (2010). A General Approach to Causal Mediation Analysis. Psychological methods 15, 309--334.

Jennes, L., and Langub, M.C. (2000). Endocrine Targets in the Brain BT - Neuroendocrinology in Physiology and Medicine (nHumana Press).

Jiang, X., Chevillet, M.A., Rauschecker, J.P., and Riesenhuber, M. (2018). Training Humans to Categorize Monkey Calls: Auditory Feature-and Category-Selective Neural Tuning Changes. Neuron 98, 405–416 e404.

Kaernbach, C. (1991). Simple adaptive testing with the weighted up-down method. Perception & psychophysics 49, 227–229.

Katsu, Y., and Iguchi, T. (2016). Cortisol. Part III: Lipophilic Hormones in Vertebrates, 533–595 –532.

Kirschbaum, C., and Hellhammer, D.H. (1989). Salivary cortisol in psychobiological research: an overview. Neuropsychobiology 22, 150–169.

Kirschbaum, C., and Hellhammer, D.H. (1994). Salivary cortisol in psychoneuroendocrine research: Recent developments and applications. Psychoneuroendocrinology 19, 313–333.

Kirschbaum, C., Prussner, J.C., Stone, A.A., Federenko, I., Gaab, J., Lintz, D., Schommer, N., and Hellhammer, D.H. (1995). Persistent high cortisol responses to repeated psychological stress in a subpopulation of healthy men. Psychosom Med 57, 468–474.

Lee, B.K., Glass, T.A., McAtee, M.J., and Wand, G.S. (2007). Associations of Salivary Cortisol With Cognitive Function in the Baltimore Memory Study. Arch Gen Psychiatry 64, 810–818.

Lezak, M.D. (1995). Neuropsychological assessment.

Linden, J.F. (2017). Timing Is Everything: Corticothalamic Mechanisms for Active Listening. Neuron 95, 3–5.

Lüdecke, D. (2020). Extracting, Computing and Exploring the Parameters of Statistical Models using R. Journal of Open Source Software 5, 2445.

Luke, S.G. (2017). Evaluating significance in linear mixed-effects models in R. Behavior Research Methods 49, 1494--1502.

Lupien, S.J., and Lepage, M. (2001). Stress, memory, and the hippocampus: can’t live with it, can’t live without it. Behavioural Brain Research 127, 137–158.

Meltser, I., Cederroth, C.R., Basinou, V., Savelyev, S., Lundkvist, G.S., and Canlon, B. (2014). TrkB-Mediated Protection against Circadian Sensitivity to Noise Trauma in the Murine Cochlea. Current Biology 24, 658–663.

Nakagawa, S., Johnson, P.C.D., and Schielzeth, H. (2017). The coefficient of determination R(2) and intra-class correlation coefficient from generalized linear mixed-effects models revisited and expanded. Journal of the Royal Society Interface 14, 20170213.

Park, H.D., Correia, S., Ducorps, A., and Tallon-Baudry, C. (2014). Spontaneous fluctuations in neural responses to heartbeats predict visual detection. Nature neuroscience 17, 612–618.

Park, J.S., Cederroth, C.R., Basinou, V., Meltser, I., Lundkvist, G., and Canlon, B. (2016). Identification of a Circadian Clock in the Inferior Colliculus and Its Dysregulation by Noise Exposure. J Neurosci 36, 5509–5519.

Prins, N., and Kingdom, F.A.A. (2018). Applying the Model-Comparison Approach to Test Specific Research Hypotheses in Psychophysical Research Using the Palamedes Toolbox. Front Psychol 9, 1250.

Pruessner, J.C., Wolf, O.T., Hellhammer, D.H., Buske-Kirschbaum, A., Auer, K.V., Jobst, S., and Kirschbaum, C. (1997). Free cortisol levels after awakening: a reliable biological marker for the assessment of adrenocortical activity. Life Sciences 61, 2539–2549.

Rarey, K.E., and Curtis, L.M. (1996). Receptors for Glucocorticoids in the Human Inner Ear. Otolaryngology–Head and Neck Surgery 115, 38–41.

Rebollo, I., Devauchelle, A.D., Beranger, B., and Tallon-Baudry, C. (2018). Stomach-brain synchrony reveals a novel, delayed-connectivity resting-state network in humans. eLife 7.

Roberts, A.C., Robbins, T.W., Weiskrantz, L., and Royal Society (Great Britain). Discussion Meeting. (1998). The prefrontal cortex: executive and cognitive functions (Oxford; New York: Oxford University Press).

Schwabe, L., Bohringer, A., Chatterjee, M., and Schachinger, H. (2008). Effects of pre-learning stress on memory for neutral, positive and negative words: Different roles of cortisol and autonomic arousal. Neurobiol Learn Mem 90, 44–53.

Singer, W., Kasini, K., Manthey, M., Eckert, P., Armbruster, P., Vogt, M.A., Jaumann, M., Dotta, M., Yamahara, K., Harasztosi, C., et al. (2018). The glucocorticoid antagonist mifepristone attenuates sound-induced long-term deficits in auditory nerve response and central auditory processing in female rats. FASEB J 32, 3005–3019.

Spencer, R.L., and Deak, T. (2017). A users guide to HPA axis research. Physiol Behav 178, 43–65.

Spiga, F., Walker, J.J., Terry, J.R., and Lightman, S.L. (2014). HPA axis-rhythms. Comprehensive Physiology 4, 1273–1298.

Stawski, R.S., Almeida, D.M., Lachman, M.E., Tun, P.A., Rosnick, C.B., and Seeman, T. (2011). Associations between cognitive function and naturally occurring daily cortisol during middle adulthood: timing is everything. Journals of Gerontology Series B: Psychological Sciences and Social Sciences 66, 71– 81.

Stolz, G., Aschoff, J.C., Aschoff, J., and Born, J. (1987). Circadian variation in the visual evoked potential (VEP). Electroencephalogr Clin Neurophysiol Suppl 40, 279–283.

Tassi, P., and Pins, D. (2009). Diurnal Rhythmicity for Visual Sensitivity in Humans? Chronobiology International 14, 35–48.

Tingley, D., Yamamoto, T., Hirose, K., Keele, L., and Imai, K. (2014). mediation: R Package for Causal Mediation Analysis. Journal of Statistical Software 59.

Trune, D.R., and Canlon, B. (2012). Corticosteroid therapy for hearing and balance disorders. Anat Rec (Hoboken) 295, 1928–1943.

Tune, S., Wostmann, M., and Obleser, J. (2018). Probing the limits of alpha power lateralisation as a neural marker of selective attention in middle-aged and older listeners. Eur J Neurosci 48, 2537–2550.

Wagenmakers, E.J. (2007). A practical solution to the pervasive problems of p values. Psychonomic Bulletin & Review 14, 779–804.

Waschke, L., Tune, S., and Obleser, J. (2019). Local cortical desynchronization and pupil-linked arousal differentially shape brain states for optimal sensory performance. eLife 8.

Wilhelm, I., Born, J., Kudielka, B.M., Schlotz, W., and Wust, S. (2007). Is the cortisol awakening rise a response to awakening? Psychoneuroendocrinology 32, 358–366.

Wolf, O.T., Convit, A., Thorn, E., and Leon, M.J. (2002). Salivary cortisol day profiles in elderly with mild cognitive impairment. Psychoneuroendocrinology 27, 777–789.

Yerkes, R.M., and Dodson, J.D. (1908). The Relation of Strength of Stimulus to rapidity of habit-formation. Journal of Comparative Neurology and Psychology 18, 459–482.

Zekveld, A.A., van Scheepen, J.A.M., Versfeld, N.J., Veerman, E.C.I., and Kramer, S.E. (2019). Please try harder! The influence of hearing status and evaluative feedback during listening on the pupil dilation response, saliva-cortisol and saliva alpha-amylase levels. Hearing Research 381, 107768.

