## Supplement for "Circadian fluctuations in glucocorticoid level predict perceptual discrimination sensitivity"

#### Supplemental information
#### for

### Circadian fluctuations in glucocorticoid level predict perceptual discrimination sensitivity

Jonas Obleser [1,3*],
Jens Kreitewolf [1,4,5],
Ricarda Vielhauer [1],
Fanny Lindner [1],
Carolin David [1],
Henrik Oster [2,3 §]
& Sarah Tune [1,3 §]

1 – Department of Psychology, University of Lübeck, 23562 Lübeck, Germany
2 – Institute of Neurobiology, University of Lübeck, 23562 Lübeck, Germany
3 – Center for Brain, Behavior, and Metabolism, University of Lübeck, 23562 Lübeck, Germany
4 – Department of Psychology, McGill University, Montréal, Canada
5 – Department of Mathematics and Statistics, McGill University, Montréal, Canada

§ shared senior authorship

#### Supplemental figures

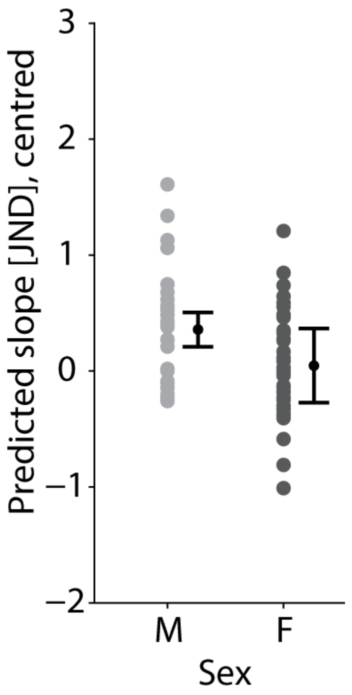


**Figure S1. Perceptual discrimination sensitivity as a function of sex.**
Coloured dots (light grey, male (M); dark grey, female (F) cohort) show single-subject (N=68) predicted slope values based on the best-fitting linear mixed-effects model. Black dots represent the fixed-effect group-level prediction and 95% CI.

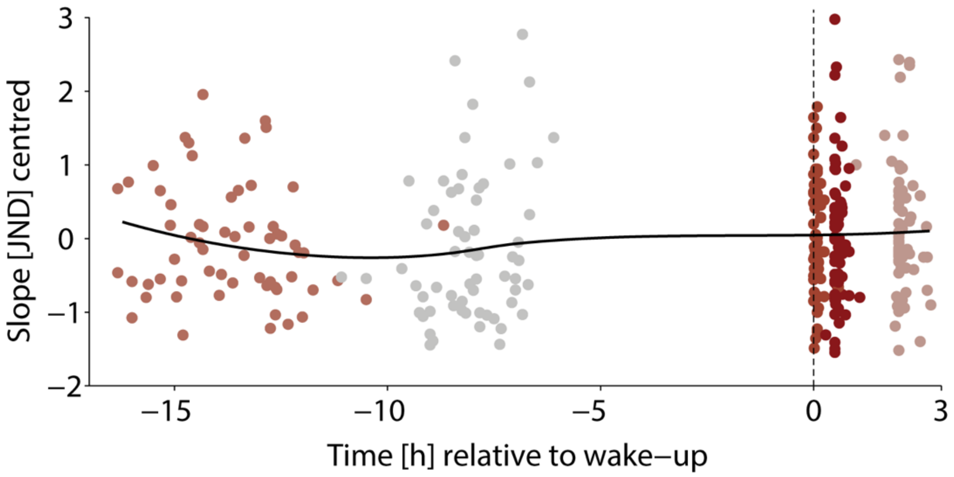


**Figure S2. Perceptual discrimination sensitivity does not depend on the time of day.**

Change in individual perceptual sensitivity across five experimental session. Slope values are mean-centred across all N=68 participants. Sessions are grouped by colour and aligned by wake-up time (dashed vertical line). Black curve shows LOESS regression of time.

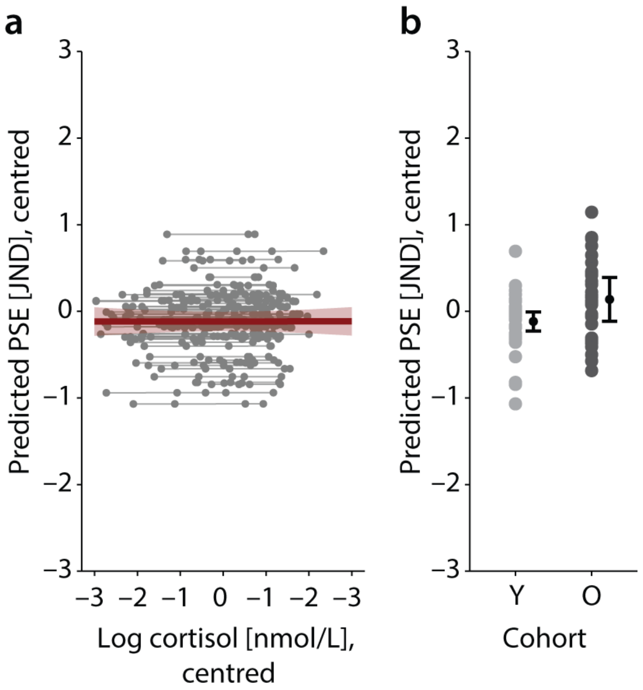


**Figure S3. Cortisol dynamics do not impact response bias.**

**(a)** Response bias (operationalised by the point-of-subjective-equality; PSE) as predicted by cortisol. Predicted non-significant group-level fixed-effect (green slope) with 95% confidence interval (CI) error band is shown along with the estimated subject-specific (N=68) random slopes (thin grey lines) and single-subject, single-session predictions (grey dots). Note that subject-specific random slopes did not improve the model fit and were added for illustrative purposes only.

**(b)** Change in response bias as a function of age cohort. Coloured dots (light grey, young (Y) cohort; dark grey, older (O) cohort) show single-subject (N=68) predicted PSE values based on the best-fitting linear mixed-effects model. Black dots represent the fixed-effect group-level prediction and 95% CI.

#### Supplemental tables

**Table S1: Predicting Cortisol**

| | Cortisol [log nmol/L; z-scored] | | | | | |
|---|---|---|---|---|---|---|
| *Predictors* | *Estimates* | *std. Error* | *CI* | *t* | *p* | *df* |
| Intercept | -0.01 | 0.24 | -0.81 – 0.80 | -0.03 | 0.979 | 2.82 |
| Time rel. to wake-up [h; z-scored] | 0.57 | 0.23 | -0.06 – 1.20 | 2.43 | 0.067 | 4.31 |
| Time squared [z] | 0.11 | 0.11 | -0.12 – 0.34 | 1.01 | 0.323 | 20.94 |
| Time cubic [z] | -0.15 | 0.07 | -0.28 – -0.01 | -2.13 | **0.035** | 128.17 |
| Sleep duration [h; z-scored] | -0.16 | 0.04 | -0.24 – -0.07 | -3.57 | **0.001** | 77.48 |
| **Random Effects** | | | | | | |
| $\sigma^2$ | 0.24 | | | | | |
| $\tau_{00\ subj}$ | 0.06 | | | | | |
| $\tau_{00\ session}$ | 0.29 | | | | | |
| ICC | 0.59 | | | | | |
| $N_{subj}$ | 68 | | | | | |
| $N_{session}$ | 5 | | | | | |
| Observations | 318 | | | | | |
| Marginal $R^2$ / Conditional $R^2$ | 0.388 / 0.750 | | | | | |
| AIC | 553.699 | | | | | |

**Table S2: Predicting Perceptual Sensitivity**

| | Perceptual Sensitivity [JND; z-scored] | | | | | |
|---|---|---|---|---|---|---|
| *Predictors* | *Estimates* | *std. Error* | *CI* | *t-value* | *p* | *df* |
| Intercept | -0.024 | 0.088 | -0.199 – 0.152 | -0.270 | 0.7883 | 64.668 |
| Cortisol [log nmol/L; z-scored] | 0.130 | 0.044 | 0.043 – 0.217 | 2.940 | **0.0036** | 267.356 |
| Age Cohort (Old) | -0.516 | 0.178 | -0.872 – -0.160 | -2.896 | **0.0051** | 65.476 |
| Sex (Female) | -0.360 | 0.180 | -0.718 – -0.001 | -2.004 | **0.0492** | 65.098 |
| **Random Effects** | | | | | | |
| $\sigma^2$ | 0.55 | | | | | |
| $\tau_{00\ subj}$ | 0.39 | | | | | |
| ICC | 0.42 | | | | | |
| $N_{subj}$ | 68 | | | | | |
| Observations | 318 | | | | | |
| Marginal $R^2$ / Conditional $R^2$ | 0.089 / 0.470 | | | | | |
| AIC | 829.669 | | | | | |

**Table S3: Predicting Perceptual Sensitivity from State- and Trait-level Cortisol**

| | **Perceptual Sensitivity [JND; z-scored]** | | | | | |
|---|---|---|---|---|---|---|
| *Predictors* | *Estimates* | *std. Error* | *CI* | *t-value* | *p* | *df* |
| Intercept | -0.024 | 0.088 | -0.201 – 0.153 | -0.269 | 0.7888 | 63.752 |
| Cortisol within-subject effect [log nmol/L; z-scored] | 0.120 | 0.042 | 0.038 – 0.202 | 2.885 | **0.0043** | 249.107 |
| Cortisol between-subject effect [log nmol/L; z-scored] | 0.051 | 0.088 | -0.124 – 0.226 | 0.580 | 0.5641 | 64.762 |
| Age Cohort (Old) | -0.516 | 0.180 | -0.875 – -0.158 | -2.875 | **0.0055** | 64.493 |
| Sex (Female) | -0.360 | 0.183 | -0.726 – 0.006 | -1.967 | 0.0535 | 63.736 |
| **Random Effects** | | | | | | |
| $\sigma^2$ | 0.55 | | | | | |
| $\tau_{00\ subj}$ | 0.40 | | | | | |
| ICC | 0.42 | | | | | |
| $N_{subj}$ | 68 | | | | | |
| Observations | 318 | | | | | |
| Marginal $R^2$ / Conditional $R^2$ | 0.088 / 0.474 | | | | | |
| AIC | 834.828 | | | | | |

**Table S4: Predicting Response Bias**

| | **PSE [point of subjective equality; z-scored]** | | | | | |
|---|---|---|---|---|---|---|
| *Predictors* | *Estimates* | *std. Error* | *CI* | *t-value* | *p* | *df* |
| Intercept | 0.019 | 0.099 | -0.178 – 0.215 | 0.189 | 0.8509 | 66.036 |
| Cortisol [log nmol/L; z-scored] | 0.001 | 0.041 | -0.080 – 0.082 | 0.022 | 0.9823 | 261.837 |
| Age Cohort (Old) | 0.445 | 0.197 | 0.052 – 0.839 | 2.260 | **0.0271** | 66.035 |
| **Random Effects** | | | | | | |
| $\sigma^2$ | 0.47 | | | | | |
| $\tau_{00\ subj}$ | 0.55 | | | | | |
| ICC | 0.54 | | | | | |
| $N_{subj}$ | 68 | | | | | |
| Observations | 318 | | | | | |
| Marginal $R^2$ / Conditional $R^2$ | 0.046 / 0.564 | | | | | |
| AIC | 803.468 | | | | | |

**Table S5: Overview of prescription medication reported by cohort of middle-aged and older participants**

| Purpose | Medication | N |
|---|---|---|
| Asthma medication | Alvesco (Ciclesonid) | 1 |
| | Amlodipin | 5 |
| | Atacant | 3 |
| | Bisoporol | 5 |
| | Candesartan | 1 |
| | Enalapril | 1 |
| Hypertension | Hydrochlorothiazide | 1 |
| | Metopropol | 1 |
| | Ramipril | 4 |
| | Valsartan | 1 |
| | Vocado40 | 1 |
| Lipid-lowering medication | Simvastatin | 1 |
| Non-steroidal anti-inflammatory medication | Aspirin | 1 |
| | Naproxen | 1 |
| | Salofalk | 1 |
